# Defense hormones modulate root microbiome diversity and composition in tomato

**DOI:** 10.1101/656769

**Authors:** Elizabeth French, Manoj Ghaste, Joshua R. Widhalm, Anjali S. Iyer-Pascuzzi

## Abstract

Plant roots live closely associated with soil microbes. Understanding how roots defend against pathogenic microbes while associating with non-pathogenic microbes is critical to using root microbiomes for successful outcomes. The aim of this study was to determine the contribution of the plant defense hormones ethylene (ET), jasmonic acid (JA), and salicylic acid (SA) to the tomato root microbiome. We used 16S rRNA amplicon sequencing to examine the root microbiome of four tomato genotypes defective in three hormone pathways. We found that ET and SA pathways, but not JA, were critical for root endosphere microbial alpha (α)-diversity. Root endosphere communities of *ACD,* deficient in a precursor of ET, and the SA-deficient *NahG* transgenic, had significantly less α-diversity than wild-type tomatoes. *NahG* and *ACD* root endosphere enrichment profiles were similar and driven by two taxa. In *NahG, ACD,* and in a panel of 24 wild-type tomatoes, the abundance of these two taxa was correlated with lower root endosphere α-diversity. The abundance of these taxa was also moderately negatively correlated with the level of SA in roots. Our results suggest that SA and ET pathways contribute to tomato root endosphere diversity by modulating the abundance of specific bacterial taxa.

## INTRODUCTION

Plant roots are intimately connected to a diversity of microbes in the soil. Some soil microbes are root pathogens and cause destructive diseases in their hosts. Others are beneficial root colonizers and contribute to plant health and function through enhancing root growth and nutrient acquisition, promoting defense responses, or augmenting abiotic stress tolerance (reviewed in (Berendsen, Pieterse & Bakker 2012; Bulgarelli, Schlaeppi, Spaepen, Ver Loren van Themaat & Schulze-Lefert 2013; Hacquard *et al.* 2015). Optimizing host selection of root-associated beneficial microbial communities holds promise for sustainably increasing crop production.

Root microbiome recruitment and assembly is determined by both the available microbes in the soil and host genotype (Lundberg *et al.* 2012; Peiffer *et al.* 2013; Schlaeppi, Dombrowski, Oter, Ver Loren van Themaat & Schulze-Lefert 2014; Haney, Samuel, Bush & Ausubel 2015; Shenton, Iwamoto, Kurata & Ikeo 2016; Pérez-Jaramillo *et al.* 2017; Fitzpatrick *et al.* 2018). The host traits that shape and select for the root microbiome are not well understood, but include root exudates, and plant hormones such as ethylene (ET), salicylic acid (SA), and jasmonic acid (JA) (Long, Sonntag, Schmidt & Baldwin 2010; Doornbos, Geraats, Kuramae, Van Loon & Bakker 2011; Carvalhais *et al.* 2013, 2015; Lebeis *et al.* 2015; Liu, Carvalhais, Schenk & Dennis 2017). The plant root microbiome can be divided into different compartments, including the rhizosphere, which is a thin layer of soil surrounding the root, and the endosphere, which is the interior of the root (Bulgarelli *et al.* 2012). Recent studies suggest that the role of plant hormones in root microbiome selection differs among species, compartment (endosphere or rhizosphere), and hormone levels (deficiency or excess) (Lebeis *et al.* 2015; Liu *et al.* 2017). For example, in Arabidopsis, the SA biosynthesis mutant *sid2* was not different than wild-type plants in root endophytic Shannon diversity (Lebeis *et al.* 2015), but the SA accumulation mutant *cpr5* had lower root endophytic Shannon diversity than wild-type plants (Lebeis *et al.* 2015).

Most work examining the role of plant hormones has been done in Arabidopsis, but other plant species can differ in how hormone pathways alter microbiomes compared to Arabidopsis (Doornbos *et al.* 2011; Santhanam, Groten, Meldau & Baldwin 2014; Liu *et al.* 2017). For example, foliar methyl jasmonate (MeJA) treatment had no effect on rhizosphere microbiome community structure or abundance in Arabidopsis (Doornbos *et al.* 2011), but in wheat, aboveground treatment with MeJA significantly reduced the Shannon diversity, richness and composition of the root endosphere, but had no impact on the rhizosphere (Liu *et al.* 2017). However, in *Nicotiana attenuata,* root associated communities of plants deficient in jasmonate biosynthesis were not different from those of control plants (Santhanam *et al.* 2014). These studies suggest that the impact of plant hormones on the root microbiome may be species-dependent. Understanding how plant hormones impact the root microbiome in crops is critical for our ability to manipulate plant-microbiome interactions for increased plant production.

Tomato is the second most important vegetable crop globally (in terms of tons produced) (FAO). Here, we hypothesized that differences in ET, SA, and JA pathways would alter tomato root microbiome structure and assembly. The role of these hormones in root microbiome assembly is particularly intriguing because these hormones are also required to defend against pathogenic microbial taxa. Given differences between tomato and Arabidopsis in ethylene regulation (Klee 1991), and JA biosynthesis, signaling (Sun, Jiang & Li 2011) and response to pathogens (Di, Gomila & Takken 2017), we also hypothesized that the role of defense hormones ET, JA, and SA in shaping the root microbiome may differ between tomato and Arabidopsis.

To test our hypotheses, we used four tomato mutants defective in ET, JA and SA hormone pathways and a panel of wild-type tomatoes to explore the function of defense hormones in tomato root microbiome structure. We find that the ET and SA pathways modulate tomato root endosphere microbiome α-diversity. Rhizosphere α-diversity was altered by the ET pathway, but was not impacted by other pathways. Additionally, we find that deficiencies in ET and SA pathways lead to an enrichment of two bacterial taxa in the root endosphere. Examination of the root endosphere microbiome in a panel of 24 wild-type tomatoes showed that the abundance of these two bacterial taxa is negatively correlated with root endosphere α-diversity. We propose that in tomato, ET and SA pathways function to modulate the abundance of specific bacterial taxa in the root endosphere.

## MATERIALS AND METHODS

### Soil mix

Soil mix was prepared by hand-mixing autoclave-sterile potting mix and field soil in a 2:1 ratio by volume. The field soil was a sandy loam collected from the top 10 cm of a conventional agricultural field at Throckmorton Purdue Agricultural Center (40.2° N, 86.9° W) in three batches from April – June 2017, ground, sieved to 4 mm, air dried at 27 °C to a constant weight, and mixed to homogenize the three batches. Potting mix was Fafard germination mix, custom blend with 56.69% spaghnum peat moss, composted bark, perlite, vermiculite, dolomite lime, wetting agent and 0.001% silicon dioxide (SKU code 8269028, lot Q17.05). The potting mix was autoclaved for 30 min at 122.8 °C. Samples of field soil and potting mix/field soil mixture were sent to A&L Great Lakes Laboratories for nutrient characterization (Table S1). Three technical replicates were performed for each soil sample.

### Plant genotypes

To test the effect of defense hormones on the root microbiome, the following genotypes were used: the *NahG* transgenic constitutively degrades SA, and is in the *Solanum lycopersicum* Moneymaker (MM) background (Brading, Hammond-Kosack, Parr & Jones 2000). *def1* is a JA biosynthesis mutant in the background of *S. lycopersicum* Castlemart II (CMII) (Howe 1996). *Neverripe* (*Nr*) is a spontaneous ethylene perception and signaling mutant in the near isogenic *S. lycopersicum* cultivar Pearson (Lanahan 1994). Although *Nr* plants are deficient in ET perception, they produce the ethylene precursor 1-aminocyclopropane-1-carboxylic acid (ACC) and ET, and are not impaired in any step of ethylene biosynthesis (Lanahan 1994). *ACD* is a transgenic which constitutively degrades ACC. *ACD* plants have low levels of both ACC and ET (Lanahan 1994). *ACD* is in the background of the *S. lycopersicum* cultivar UC82B (Klee 1991). Although *NahG* and *ACD* are transgenic plants, we refer to them as ‘mutants’ for simplicity. Mutant lines and backgrounds are listed in Table S2.

To test whether two bacterial taxa enriched in *NahG* and *ACD* root endospheres were generally associated with lower α-diversity, we examined a panel of 20 recombinant inbred lines (RILs) derived from the *S. lycopersicum* cv. Hawaii7996 (H7996) and the *S. pimpinellifolium* West Virginia700 (WV). To control for species effects, we included another *S. lycopersicum* (cv. Bonny Best) and another *S. pimpinellifolium* (LA2093). Parents and RILs are listed in Table S3. Microbiome analyses of hormone mutants, wild-type background, RILs and parents was performed together as described below.

### Plant growth and harvest

Seeds were surface sterilized by incubating with gentle rocking in 50% bleach for ten minutes and then rinsing 5-6 times in sterile ddH_2_O. Seeds were then stored at 4 °C overnight. Surface-sterile seeds were planted in randomized complete blocks in 36-pot flats with 1 unplanted cell per block to represent the bulk soil control. Three seeds were planted to each pot and thinned to one plant after germination. Eight full biological replicate blocks were planted to account for any issues with germination. Flats were fertilized once per week with 500 mL of 150 ppm Nitrogen standard MiracleGro fertilizer. Plants were grown in a light and temperature controlled greenhouse (temperature setting 75-84°F). Lights operated on a 16 hour on, 8 hour off long day cycle.

Four blocks were harvested for sequencing after seedlings reached 4-leaf stage (approximately 2.5 weeks). Rhizosphere and endosphere samples were collected from each genotype except RILs, for which only endosphere samples were collected (see Tables S2 and S3 for all genotypes). For rhizosphere sampling, roots were removed from the pot and excess soil was removed gently under aseptic conditions until only soil within 1 mm from the root surface remained (Lundberg *et al.* 2012). Roots were then placed into 15 mL conical tubes containing sterile 1X PBS, shaken manually, and then placed in a new 15 mL conical tube for surface sterilization. Conical tubes containing the rhizosphere soil were spun at 5000 rpm for 5 min. Excess liquid was decanted, and soil pellets were resuspended and transferred to a 1.5 mL sterile Eppendorf tube. The Eppendorf tube was spun at max speed for 10 min, supernatant decanted, rhizosphere soil frozen in liquid nitrogen (LN_2_) and stored at -80 °C until DNA extraction. These samples were designated the rhizosphere compartment samples.

Roots were cleaned by performing an additional rinse to remove any remaining soil. Then roots were surface sterilized by incubating in 5% bleach with gentle shaking for 2 min, then rinsed 3 times in sterile ddH_2_O before freezing in LN_2_ and storing at -80 °C until DNA extraction. These samples were designated the endosphere compartment samples.

### DNA extraction and library preparation

Frozen roots were ground under liquid nitrogen (LN_2_) in sterile mortars and pestles before DNA extraction. For the hormone mutant and respective wild-type backgrounds, each root sample weighed approximately 0.23 ± 0.02 g. For the RILs, H7996, BB, WV, and LA2093, each root sample weight approximately 0.21 ± 0.02 g. DNA was extracted from all samples using Norgen Soil DNA (Norgen Biotek Corp, Canada) extraction kits. DNA concentration and purity were measured by Nanodrop3000. Library preparation was performed using the Illumina 16S Metagenomics Sequencing Library Preparation protocol according to the manufacturer’s instructions with slight modifications. Two step PCR was performed to amplify the V5 through V7 region of the 16S rRNA gene with the chloroplast excluding primer pair 799F-1193R (Chelius & Triplett 2001; Beckers *et al.* 2016) and to add Illumina Nextera XT indices.

First, all DNA samples were diluted to 5 ng/µL. One negative water control and one mock community DNA control (ZymoBIOMICS Microbial Community Standard, Zymo Research, Irvine, CA, USA) were included on each plate. 25 µL PCR reactions were performed with 2.5 µL genomic DNA, 12.5 µL 2X KAPA HiFi HotStart ReadyMix and 5 µL each of the forward and reverse primers (1 µM) in two 96 well plates. Primer sequences with adapters were as follows: 799F + Nextera adapter: 5’-TCGTCGGCAGCGTCAGATGTGTATAAGAGACAG<u>AACMGGATTAGATACCCKG</u>-3’ and 1193R + Nextera adapter: 5’-GTCTCGTGGGCTCGGAGATGTGTATAAGAGACAG<u>ACGTCATCCCCACCTTCC</u>-3’, which produces a ∼480 bp product. Underlined portions indicate the 16S primer portion. PCR cycle performed as follows: 95 °C for 3 min, [95 °C for 30 sec, 55 °C for 30 sec, 72 °C for 30 sec] x 27 cycles, 72 °C for 5 min.

After the initial PCR step, PCR products were run on 1.5% agarose gels. The ∼480 bp band was excised and gel extracted with the Invitrogen PureLink gel purification kit. This step was performed to exclude the larger mitochondrial band amplified by the 16S primers. Gel purified PCR products were used for the second PCR step to attach dual indices and sequencing adapters. For this step, 50 µl PCR reactions were performed with 5 µL purified Step 1 PCR product, 25 µL 2X KAPA HiFi HotStart ReadyMix, 10 µL of sterile ddH2O and 5 µL each of the Index 1 and Index 2 Nextera XT Primers (set A and B) in two 96 well plates. PCR cycle was as follows: 95 °C for 3 min, [95 °C for 30 sec, 55 °C for 30 sec, 72 °C for 30 sec] x 8 cycles, 72 °C for 5 min. Standard AmpureXP bead purification was performed on the Step 2 PCR products.

Step 2 PCR products were quantified at the Purdue Genomics Core by mixing 1 µL of each library into a pool and sequencing as 10% of a MiSeq paired end 250 bp run. The library sizes were estimated from the number of reads obtained from each library and used to calibrate library concentrations for the final pool. All 188 samples were multiplexed into a single pool in equivalent concentrations. The pool was run on an Agilent bioanalyzer chip to confirm library size and purity. The pool was sequenced at the Purdue Genomics Core Facility using Illumina MiSeq V2 chemistry with paired end 250 bp sequencing.

### Sequence processing

Demultiplexing was performed by the Purdue Genomics Core with Illumina software; adapter removal and primer clipping was performed with Trimmomatic (v 0.36) (Bolger, Lohse & Usadel 2014) and Cutadapt (v 1.13) (Martin 2011). All subsequent processing was performed using packages in R (v 3.5.0) and Bioconductor (v 3.7). Reads were processed through the DADA2 (v 1.8.0) pipeline by filtering and trimming based on read quality, inferring error rates, merging paired end reads, removing chimeras, and assigning taxonomy with the Silva reference database v. 132 (Callahan *et al.* 2016). Likely contaminant sequences were removed with the decontam package using negative controls to infer likely contaminants (Davis, Proctor, Holmes, Relman & Callahan 2018). Very low abundance sequences (fewer than 2 reads in 10% of the samples) were removed. Samples with α-diversity measurements more than 1.5X outside the interquartile range were considered outliers and removed. α-diversity measurements performed with the Phyloseq (v 1.24.0) package after subsampling to the smallest library size (2,569 reads) 100 times and averaging the results (McMurdie & Holmes 2013). β-diversity measurements were performed with Phyloseq and vegan (v 2.5-2) packages with reads proportionally scaled to the smallest library size (code courtesy of Denef lab tutorial -http://deneflab.github.io/MicrobeMiseq/). Normalization and differential abundance analysis were performed with DESeq2 (v 1.20.0) (Love, Huber & Anders 2014). Linear regression performed with lm() function in R. All plots were made with the ggplot2 (Wickham 2009) package and arranged in Inkscape (v 0.92.3). All code for analysis and figure generation can be found in the Purdue University Github (https://github.rcac.purdue.edu) as ‘Tomato-Root-16S-Sequencing’. Sequencing summary is listed in Table S4.

### Total SA quantification

Plants were grown in a light and temperature controlled greenhouse as above in the same soil mix and harvested at the same stage. 300 mg (±10) of fresh root tissue (ground to a powder in LN_2_) was mixed with 3 mL 80% (v/v) methanol and 80 µL of 4-chlorobenzoic acid at a concentration of 5 µg/mL, and extracted overnight by shaking (200 rpm) at 4 °C. The tubes were centrifuged at 5000 *x g* at 4 °C for 3 min, the supernatant was removed and 1500 µL of supernatant was evaporated under nitrogen till near dryness. 500 µL 2.5 N HCl was added to the pre-evaporated tubes and incubated at 80 °C for one hour followed by ethyl acetate (5 mL) partitioning. The ethyl acetate fraction was removed and dried under nitrogen and residues were reconstituted in a final volume of 200 µL of acetonitrile and used for the LC-MS analysis.

LC-MS analysis was performed using an Agilent 1290 Infinity II UHPLC system (Palo Alto, CA, USA) coupled with 6135 single quadrupole mass spectrometer equipped with a jet stream technology electrospray ionization (ESI) source. 20 µl of sample injected and separated on a Zorbax SB-C18 column (1.8-µm, 2.1×1.8mm; Agilent) connected to a Zorbax SB-C18 analytical guard column (5-µm, 4.6 × 12.5mm; Agilent), 0.1% formic acid (v/v) in LC-MS grade water (A) and 0.1% formic acid (v/v) in acetonitrile (B) were used as the mobile phases. After sample injection, the column was eluted with 20% B for 1 min, followed by a linear gradient to 95% B over 8 min with 4 min hold, at 13 min the solvents were ramped back to 5% B with final hold of 1 min, the flow rate was maintained at 300 µL min-1 and column temperature at 40 °C. The MS was operated in negative ion mode with the nozzle and capillary voltages at 2000 V and 4000 V, respectively. Sheath gas temperature was kept at 360 °C with a flow of 12 mL min-1 and drying gas temperature kept at 350 °C with the flow of 13 mL min-1. Detection the compounds SA and 4-chlorobenzoic acid was performed in selected ion monitoring (SIM) mode with ions 137 m/z (M-H)-and 155 m/z (M-H)-respectively.

The data were analyzed using ChemStation software (version C.01.08). Identification was performed based on retention times and masses of salicylic acid and 4-chlorobenzoic acid standards (Sigma). The amount of SA was calculated on the basis of its relative response factor to the internal standard 4-chlorobenzoic acid at 1000 ng/mL. Statistical significance of differences between genotypes was tested with one-way analysis of variance (ANOVA) in JMP 13, followed by Tukey’s honest significant differences post hoc tests. Values were log transformed to meet homogeneity of variance assumption.

## RESULTS

### Tomato mutants with defects in the ET and SA pathways have significantly reduced Shannon diversity and richness in the root endosphere

To determine the role of tomato defense hormones in root microbiome assembly, we used four tomato mutants and their associated wild-type backgrounds (in tomato, mutants are frequently not in the same wild-type background). We examined the root microbiome of the following: (1) *defenseless1* (*def1*), which is defective in JA accumulation, and its wild-type background Castlemart; (2) *Neverripe (Nr),* which is defective in ET perception, and its wild-type Pearson; (3) *ACD,* a transgenic that constitutively degrades the ET precursor ACC and thus is defective in ET biosynthesis, and its wild-type UC82B; (4) *NahG,* a transgenic that constitutively degrades SA and thus is defective in SA accumulation, and its wild-type background Money Maker (see Table S2 and materials and methods for genotype information). We chose to use the *NahG* transgenic as a proxy for deficiencies in the SA pathway because SA biosynthesis mutants are not available in tomato. Similarly, we chose the *ACD* transgenic because ACC biosynthesis mutants are not available in tomato. All plants were grown in a natural soil mixed with sterile potting mix (2:1 potting mix/soil) in the greenhouse (Table S1 for soil characteristics) and DNA was harvested from bulk soil, the rhizosphere and root endosphere of four replicates of each genotype.

The V5-V7 region of the 16S rRNA from these samples was sequenced by paired-end 250 bp MiSeq Illumina sequencing, resulting in 2.9 million high quality sequences after quality filtering and removal of chimera, non-target (mitochondria, chloroplast, archaea), and likely contaminant sequences (see Materials and Methods and Table S4). These sequences corresponded to 23,833 amplicon sequence variants (ASVs). We filtered for low abundance ASVs (fewer than 2 reads in less than 10% of the samples), samples with fewer than 2000 total reads, and outliers. Our final data set for analysis consisted of approximately 1.8 million reads and 901 ASVs with an average of 10,185 reads per sample (see Materials and Methods for description of sequence processing; Tables S5-7 for raw ASV counts in each sample, sample data, and taxonomy). The 901 ASVs were comprised of 16 phyla and 123 families. The average relative abundance of the bacterial families in the rhizosphere and root endosphere of each hormone mutant and its respective wild-type background is shown in Figure 1 (taxa listed in Table S8). No significant differences between mutants and their respective wild-type backgrounds were found at the phylum level. The majority of taxa in all genotypes were in the phylum Proteobacteria and Actinobacteria.

**Figure 1.**
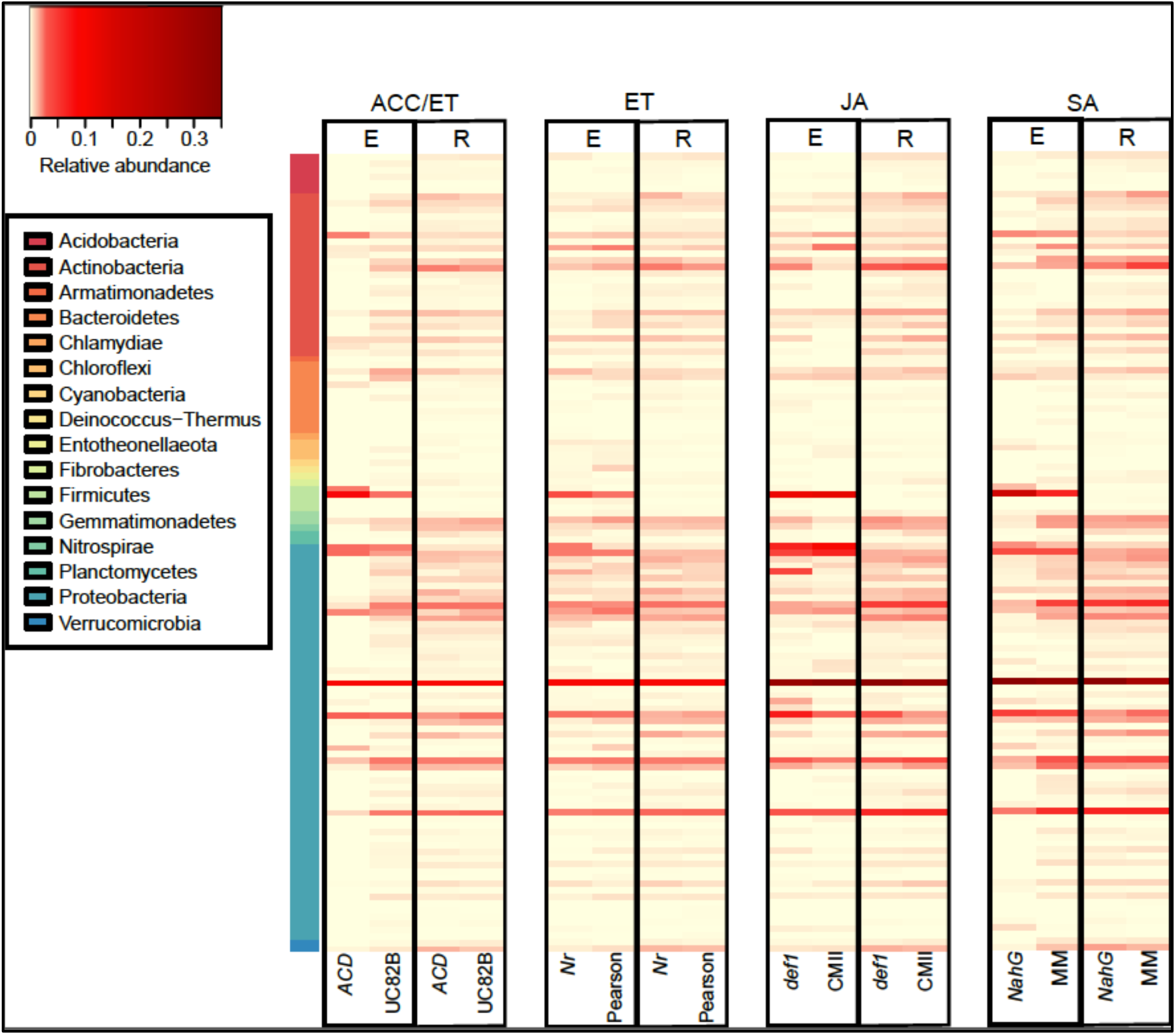
Heatmaps showing average relative abundance of 123 bacterial families across all four mutants and their respective wild-type backgrounds in the root endosphere and rhizosphere. Sidebar colors represent phyla of each family. E – endosphere. R – rhizosphere. ET – ethylene. ACC -1-aminocyclopropane-1-carboxylic acid. JA – jasmonic acid. SA – salicylic acid. *ACD* – ACC-deaminase transgenic. UC82B – *S. lycopersicum* cv. UC82B. *Nr* – *Never ripe* mutant. Pearson – *S. lycopersicum* cv. Pearson. *def1* – *defenseless1*. CMII – *S. lycopersicum* cv. Castlemart II. *NahG* – salicylate hydroxylase transgenic. MM – *S. lycopersicum* cv. Money Maker.

Bray-Curtis beta (β-)diversity patterns showed separation between the endosphere samples and the rhizosphere/bulk samples along the first axis (27.2%) and separation between the rhizosphere and bulk soil samples along the second axis (8.9%) (Figure S1) (PERMANOVA, compartment: F_(2, 154)_ = 21.96, p < 0.001). This compartmental specialization is consistent with other rhizosphere studies (Bulgarelli *et al.* 2012; Lundberg *et al.* 2012; Lebeis *et al.* 2015). Rhizosphere samples overall showed very little variation in β-diversity (Figure S1), and we chose to focus on endosphere samples for further comparisons.

*ACD* and *NahG* mutant root endosphere communities exhibited significant reductions in α-diversity (as measured by both richness and Shannon diversity) compared to their respective wild-type backgrounds (Figure 2a). No significant differences in either measure of α-diversity were observed for rhizosphere samples of *NahG*. Shannon diversity, but not richness, was significantly different between *ACD* and the rhizosphere of its wild-type background, UC82B.

**Figure 2.**
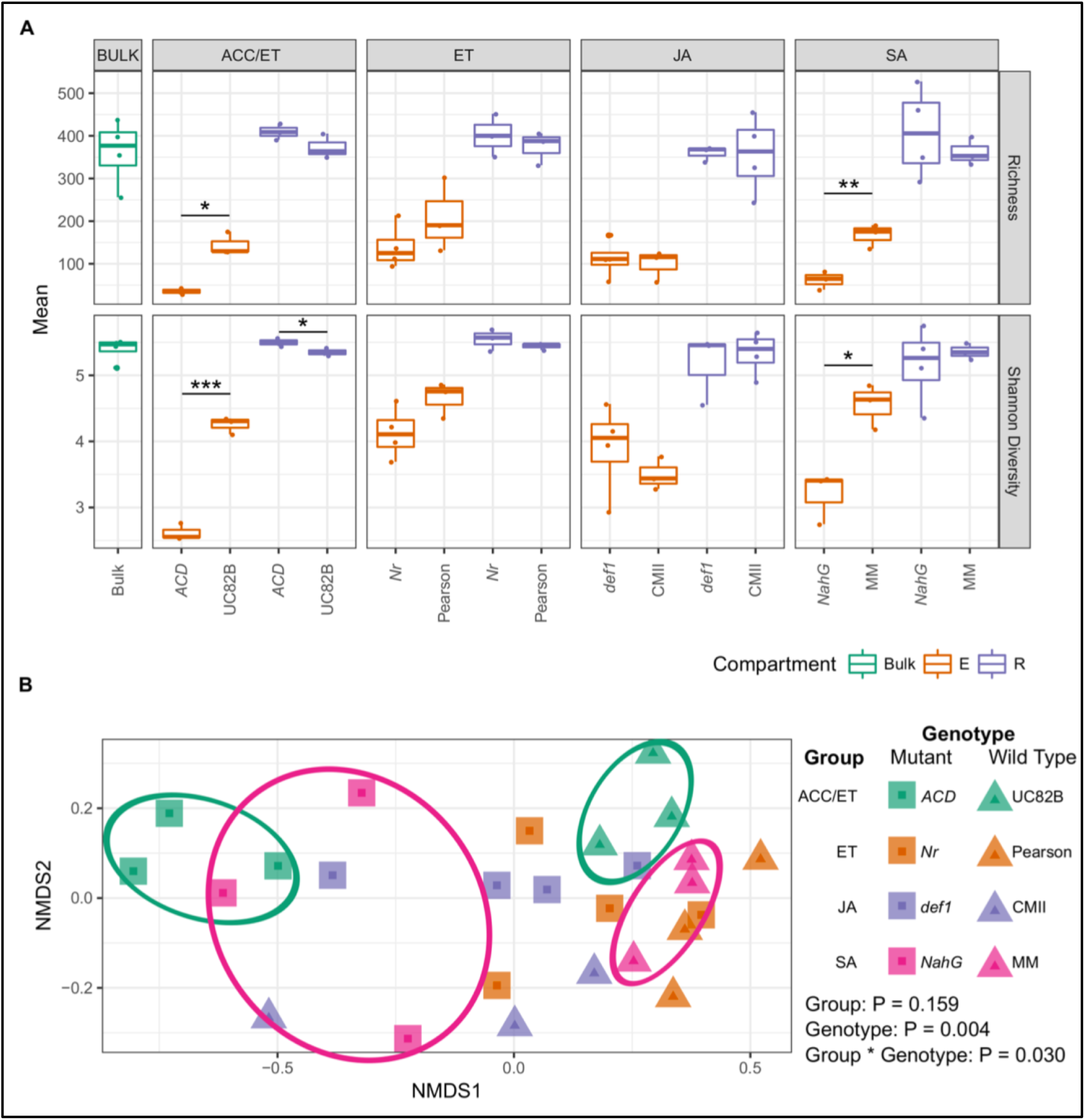
*ACD* and *NahG* mutant endospheres differ from their wild-type backgrounds in α-and β-diversity. a) Boxplots of bacterial richness and Shannon diversity of rhizospheres and endospheres of all four mutants and their respective wild-type backgrounds and bulk (unplanted) soil. Asterisks indicate significant differences by t-test, * p < 0.05. ** p < 0.01. *** p < 0.001. b) Nonmetric multidimensional scaling (NMDS) analysis of root endosphere bacterial communities of all four mutants and their wild-type backgrounds. Two-way PERMANOVA results indicated on graph. Squares indicate mutant samples, and triangles indicate wild-type samples. Colored ellipses added to emphasize group (pink = SA; green = ACC/ET). E – endosphere. R – rhizosphere. ET – ethylene. ACC -1-aminocyclopropane-1-carboxylic acid. JA – jasmonic acid. SA – salicylic acid. *ACD* – ACC-deaminase transgenic. UC82B – *S. lycopersicum* cv. UC82B. *Nr* – *Never ripe* mutant. Pearson – *S. lycopersicum* cv. Pearson. *def1* – *defenseless1*. CMII – *S. lycopersicum* cv. Castlemart II. *NahG* – salicylate hydroxylase transgenic. MM – *S. lycopersicum* cv. Money Maker.

In contrast to *ACD* and *NahG,* neither *Nr* nor *def1* were significantly different than their wild-type backgrounds for either the root endosphere or rhizosphere (Figure 2a). Notably, Shannon diversity and richness of the endosphere microbiome varied quantitatively in wild-type tomatoes, but rhizosphere diversity was less impacted by genotype (Figure 2a).

A non-metric multidimensional scaling (NMDS) analysis based on Bray-Curtis dissimilarity of root endosphere samples revealed similar patterns as the α-diversity. Both *ACD* and *NahG* separate from their wild-type backgrounds along the first axis (Figure 2b), while *def1* separated from its wild-type background along the second axis. A PERMANOVA revealed a significant effect of genotype (wild type or mutant) and a significant interaction between type of defense hormone (termed ‘Group’) and genotype (PERMANOVA Group – F_(3, 18)_ = 1.17, p = 0.159. Genotype – F_(1, 18)_ = 2.38, p = 0.004, Group*Genotype – F_(3, 18)_ = 1.46, p = 0.030) (Figure 2b).

The Bray-Curtis data in Figure 2b suggested that root endospheres of *NahG* and *ACD* may be more similar to each other than to their wild-type parents. To test this, we performed an NMDS with only *NahG, ACD* and their respective wild-types. The mutant (*NahG*/*ACD*) root endospheres separated from the wild-type genotypes along the first axis, and a PERMANOVA indicated that samples separated significantly between mutants and wild-types, but not between defense hormone groups (PERMANOVA Group – F_(1, 8)_ = 1.12, p = 0.251. Genotype– F_(1, 8)_ = 3.71, p = 0.002, Group*Genotype – F_(1, 8)_ = 1.31, p = 0.160) (Figure S2).

### Rhizosphere to endosphere differential abundance profiles reveal that ACD and NahG have additional endosphere-depleted taxa compared to their wild-type backgrounds

The root endospheres of *NahG* and *ACD* appeared to play a role in selection of the microbiome from the rhizosphere. We compared the rhizosphere to endosphere differential abundance profiles in each of the mutants and their respective wild-type controls. We used DESeq2 to determine differentially abundant ASVs between the rhizosphere and endosphere for each genotype using an adjusted p-value of 0.05 as a cutoff. Of the total 901 ASVs we identified (Table S5), 178 were differentially abundant between the rhizosphere and endosphere in at least one of the eight genotypes (4 mutant and 4 wild type –Table S4).

We categorized the 178 differentially abundant ASVs as either endosphere-enriched or endosphere-depleted (See Figure S3 for phyla distribution, Table S9 for summary, and Tables S10-17 for full results for each genotype). More endosphere-depleted ASVs were observed in mutant lines *ACD* (133) and *NahG* (109) than in their respective wild-types (UC82B = 36, MM = 32) (Figure 3a, b, the ‘down’ category is depleted; Table S6). The additional ASVs depleted in the transgenics are consistent with the reduced α-diversity observed in the endosphere of these plants. In general, most of the endosphere-depleted ASVs in either wild-type background were also found in *ACD* and *NahG* (Figure 3a-b). For example, 36 ASVs were endosphere-depleted in the wild-type background UC82B. Of these 36 ASVs, 28 (over 77%), were also depleted in *ACD*. *Nr* and *def1* had fewer depletions and shared a lower percentage of depleted ASVs with their respective wild-type backgrounds (Figure 3c-d).

**Figure 3.**
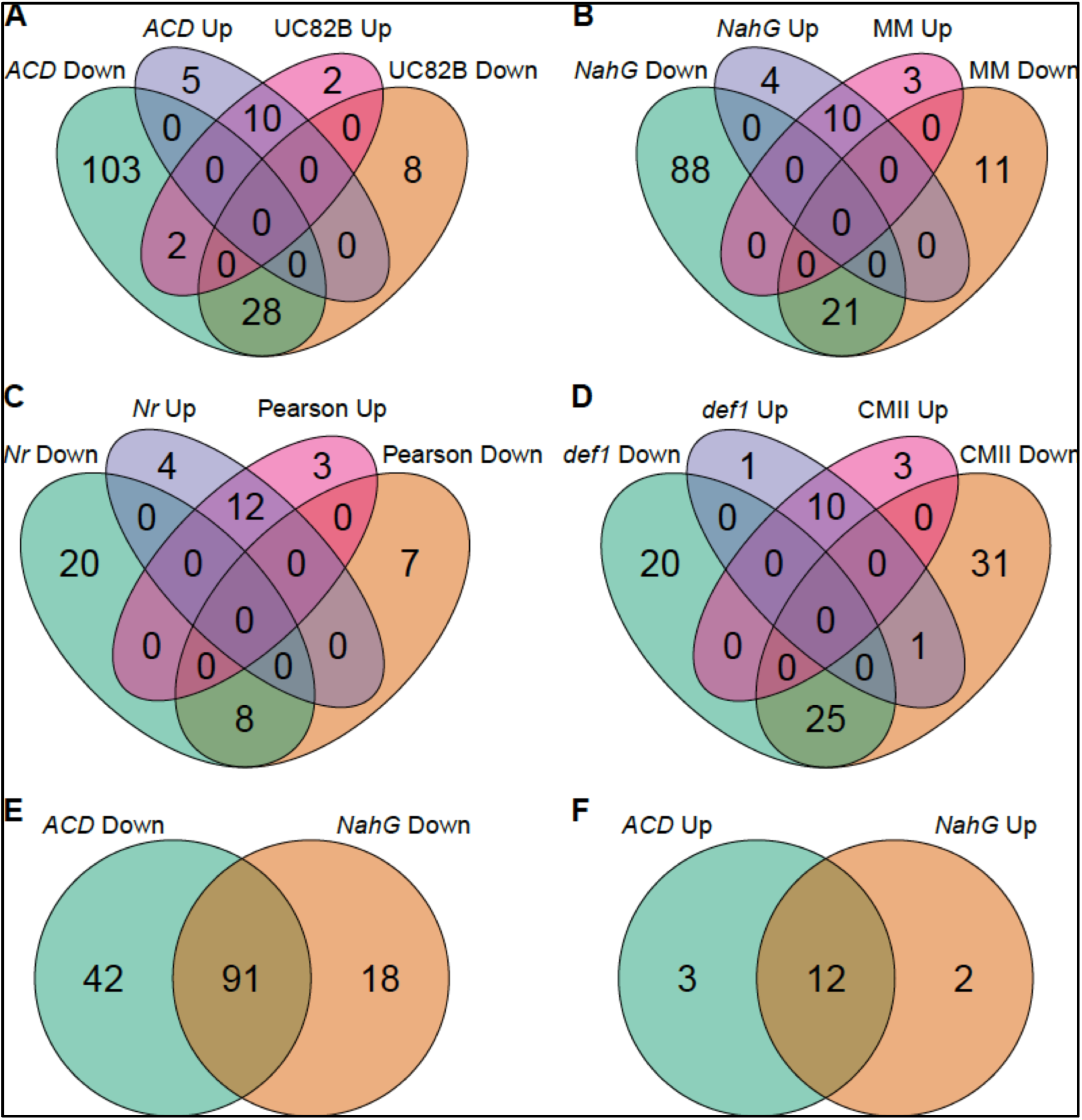
Rhizosphere to endosphere differential abundance profiles reveal additional endosphere-depletions in *NahG* and *ACD* compared to their wild-type backgrounds. Venn diagrams of enriched (Up) and depleted (Down) differentially abundant ASVs from the rhizosphere to endosphere in a) *ACD* and wild-type background UC82B, b) *NahG* and wild-type background MM, c) *Nr* and Pearson, d) *def1* and CMII. Venn diagrams comparing e) depleted ASVs and f) enriched ASVs between *NahG* and *ACD*.

Because the β-diversity pattern in Figure 2 suggested that taxa in the *ACD* and *NahG* root endospheres were more similar to each other than to those in the wild-type backgrounds, we also compared the endosphere-depleted (Figure 3e) and enriched ASVs (Figure 3f) between these two genotypes. *ACD* and *NahG* shared about 60% of depleted ASVs and 70% of enriched ASVs. Most of the shared ASVs between ACD and *NahG* belonged to the Proteobacteria, though several also belonged to Actinobacteria (Figure S4).

### Root endospheres of NahG and ACD hyperaccumulate two endosphere-enriched ASVs

Across all four hormone mutants and their wild-type backgrounds, we found a total of 27 root endosphere-enriched ASVs and 160 root endosphere-depleted ASVs, though 2 depleted ASVs had no counts in the root endosphere across all genotypes and thus were excluded from the hierarchical clustering. Some ASVs overlapped between the enriched and depleted datasets because they were enriched in one genotype and depleted in another. For simplicity, these ASVs were included in both datasets. We first calculated the relative abundances by dividing the number of reads for each ASV by the total number of reads in the sample. We then clustered the genotypes and ASVs based on the average relative abundance of enriched and depleted ASVs in the root endosphere using Bray-Curtis distance and average hierarchical clustering (Figure 4). Clustering on genotype revealed three major clusters for the enrichment profile (Figure 4a), with *NahG* and *ACD* clustered together, *Nr, def1* and CMII clustered together, and the remaining wild-type parents in a third cluster. The depletion profiles (Figure 4b) grouped in three clusters – *ACD* by itself, *NahG* and CMII together, and the remaining genotypes clustered together.

**Figure 4.**
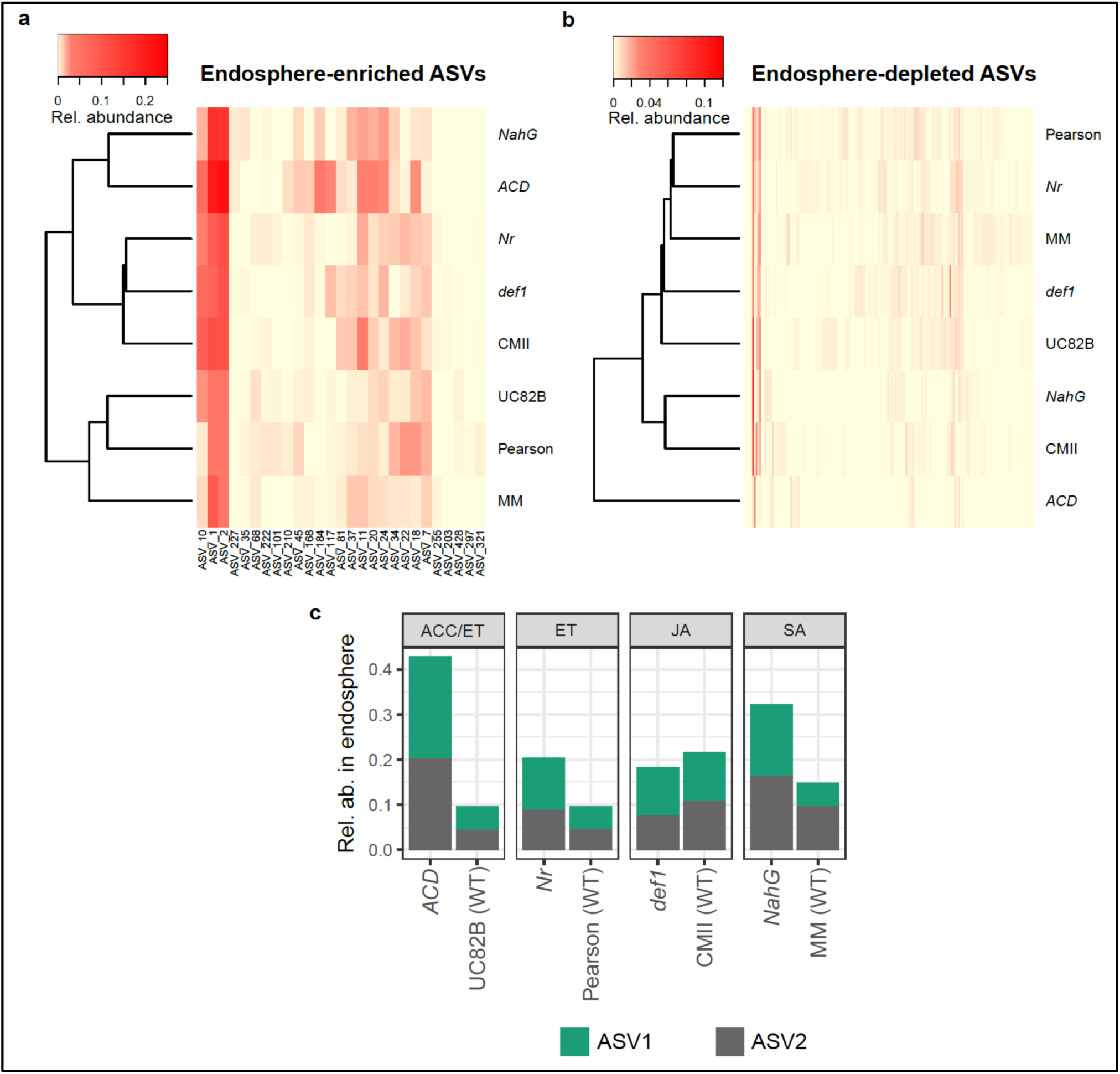
Two ASVs are highly enriched in the root endosphere of *NahG* and *ACD*. a-c) Hierarchical clustered heatmaps of average relative abundance of rhizosphere-to-endosphere a) enriched and b) depleted ASVs by genotype, c) Average relative abundance of ASV1 (*Anaerobacillus*) and ASV2 (*Delftia*) across all eight genotypes.

Examination of the endosphere-enriched ASV clusters showed that two highly abundant ASVs (termed ASV1 and ASV2) appeared to be more abundant in the endospheres of *NahG* and *ACD* compared to their wild-type backgrounds and the other genotypes (Figure 4a, c). ASV1 is classified in the genus *Anaerobacillus* (Phylum: Firmicutes) and ASV2 is in the genus *Delftia* (Phylum: Proteobacteria). Together, these two ASVs are between 30-45% of the total community of *NahG* and *ACD* but only 10-20% of the total community of the other five genotypes (Figure 4b).

### Relative abundance of NahG and ACD enriched taxa ASV1 and ASV2 is negatively correlated with Shannon diversity across wild-type tomato genotypes

*NahG* and *ACD* roots have significantly lower α-diversity than their wild-type parents (Figure 2), and they hyperaccumulate two ASVs (Figure 4). However, *ACD* and *NahG* are both transgenics that degrade either ACC or SA to other metabolites (ammonium and alpha ketobutyrate or catechol, respectively). It is possible that the lower root microbiome diversity of these lines and enrichment of ASV1/2 is due to the increased levels of ammonium, alpha ketobutyrate or catechol in these plants compared to wild-type. To examine this further, we asked whether low root endosphere diversity was correlated with enrichment of ASV1/2 in a panel of wild-type tomatoes. We sequenced the root endosphere microbiome of 20 recombinant inbred lines (RILs) derived from *S. lycopersicum* Hawaii 7996 and *S. pimpinellifolium* WV (*S. pimpinellifolium* is a wild relative to tomato), as well as Hawaii 7996, WV, *S. lycopersicum* Bonny Best and *S. pimpinellifolium* LA2093. Four replicates of each genotype were grown in the same conditions as the hormone mutants and their wild-type parents. DNA from the root endosphere was extracted and 16S rRNA amplicon sequencing was performed as above. The relative abundance of ASV1 and ASV2 in the root endosphere varied quantitatively across genotypes (Figure 5a and b), and correlated negatively with Shannon diversity (Figure 5c and d, ASV1, F_(1, 80)_ = 86.11, p = 2.482×10^-14^, R^2^ = 0.51; ASV2, F_(1, 80)_ = 85.98, p = 2.558×10^-14^, R^2^ = 0.51). Thus, similar to *NahG* and *ACD,* the abundance of ASV1/2 taxa is correlated with lower root microbiome diversity across wild-type genotypes.

**Figure 5.**
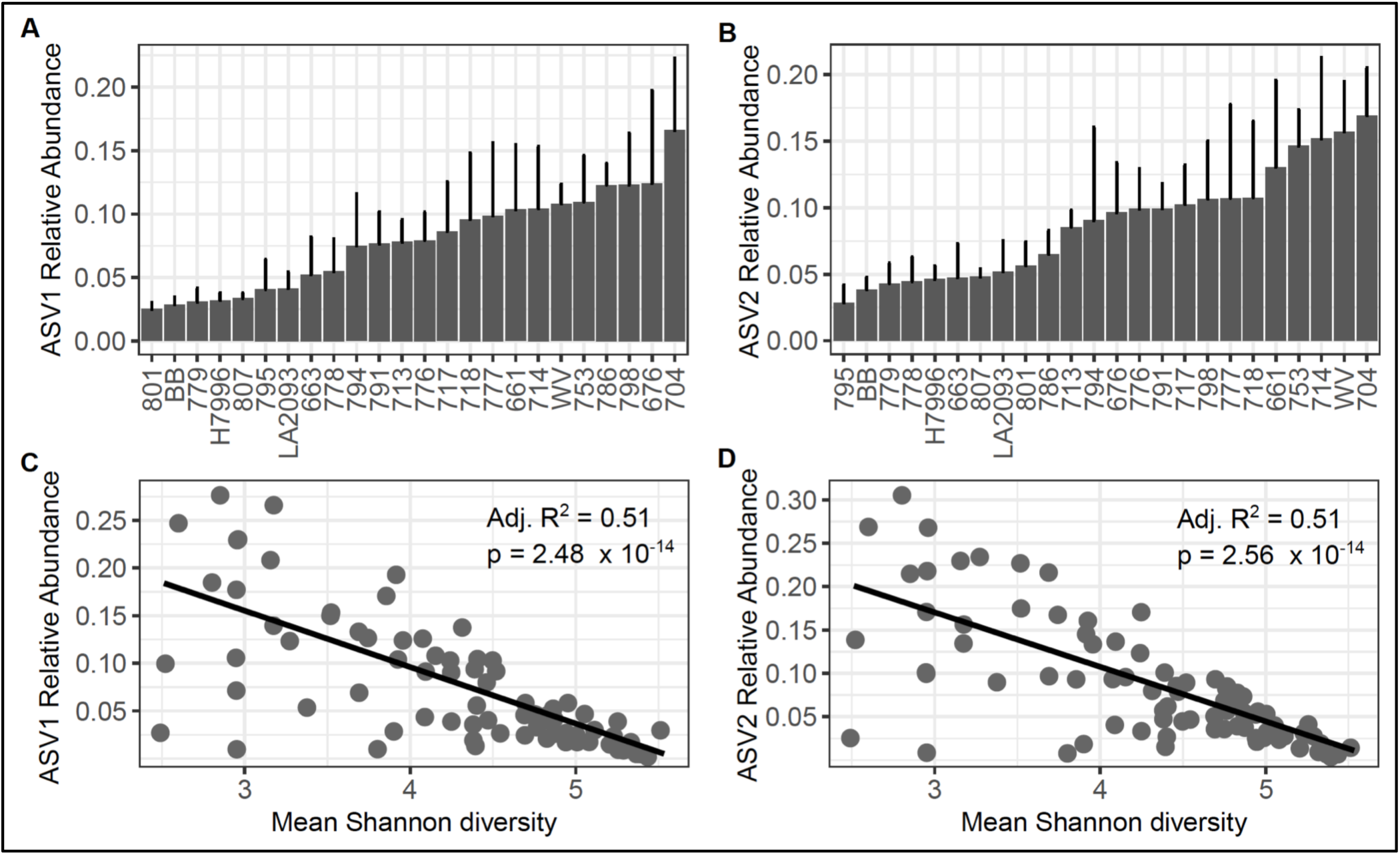
The relative abundance of ASVs 1 and 2 varies quantitatively in *S. lycopersicum* cv. H7996 and *S. pimpinellifolium* WV RILs, and correlates negatively with Shannon diversity. Relative abundance of a) ASV1 (*Anaerobacillus*) and b) ASV2 (*Delftia*) across H7996, WV, BB, LA2093 and 20 H7996 x WV RILs listed on the x-axis. Linear regression between Shannon diversity and c) ASV1 or d) ASV2. R^2^ and p values from linear regression modeling with lm() in R for each trendline represented on each plot in C and D. P values were considered significant at p < 0.05. Each data point in c) and d) is from an individual plant.

To further examine the relationship between the ET and SA pathways and the relative abundance of ASV1/2, we investigated whether there was a correlation between ET or SA levels and relative abundance of ASV1/2 in wild-type tomatoes. We measured ET and SA levels in six wild-type cultivated tomatoes and *S. pimpinellifolium* genotypes (H7996, WV, BB, UC82B, MM, LA2093). Measuring ET in roots from plants grown in soil is extremely challenging, and ET is typically measured in plant headspace (Cristescu et al. 2013). Measurements of ET in whole seedlings in cuvettes using gas chromatography found no correlation between ET levels and root endosphere diversity (data not shown). This may have been due to the organ-specificity of ET levels (Cristescu et al. 2013), or because plants used for ET measurements were grown in different conditions than those used for microbiome analyses (cuvettes vs field soil, 7 day old seedlings vs 4 week old plants). For SA measurements, plants were grown in the same field soil and greenhouse conditions as above. We found a moderate negative correlation between the relative abundance of ASV1/2 and SA levels: R^2^ = 0.399 for ASV1 and 0.477 for ASV2 (Figure 6). Thus, in wild-type tomatoes, lower root SA levels are moderately correlated with higher ASV1/2 abundance. There was no relationship between root SA level and root endosphere α-diversity (R^2^=0.08, data not shown). This suggests that SA itself does not modulate α-diversity. However, other components in the SA metabolic pathway may have a role in the process. Taken together, these data suggest that the SA pathway influences the relative abundance of specific taxa in the tomato root endosphere.

**Figure 6.**
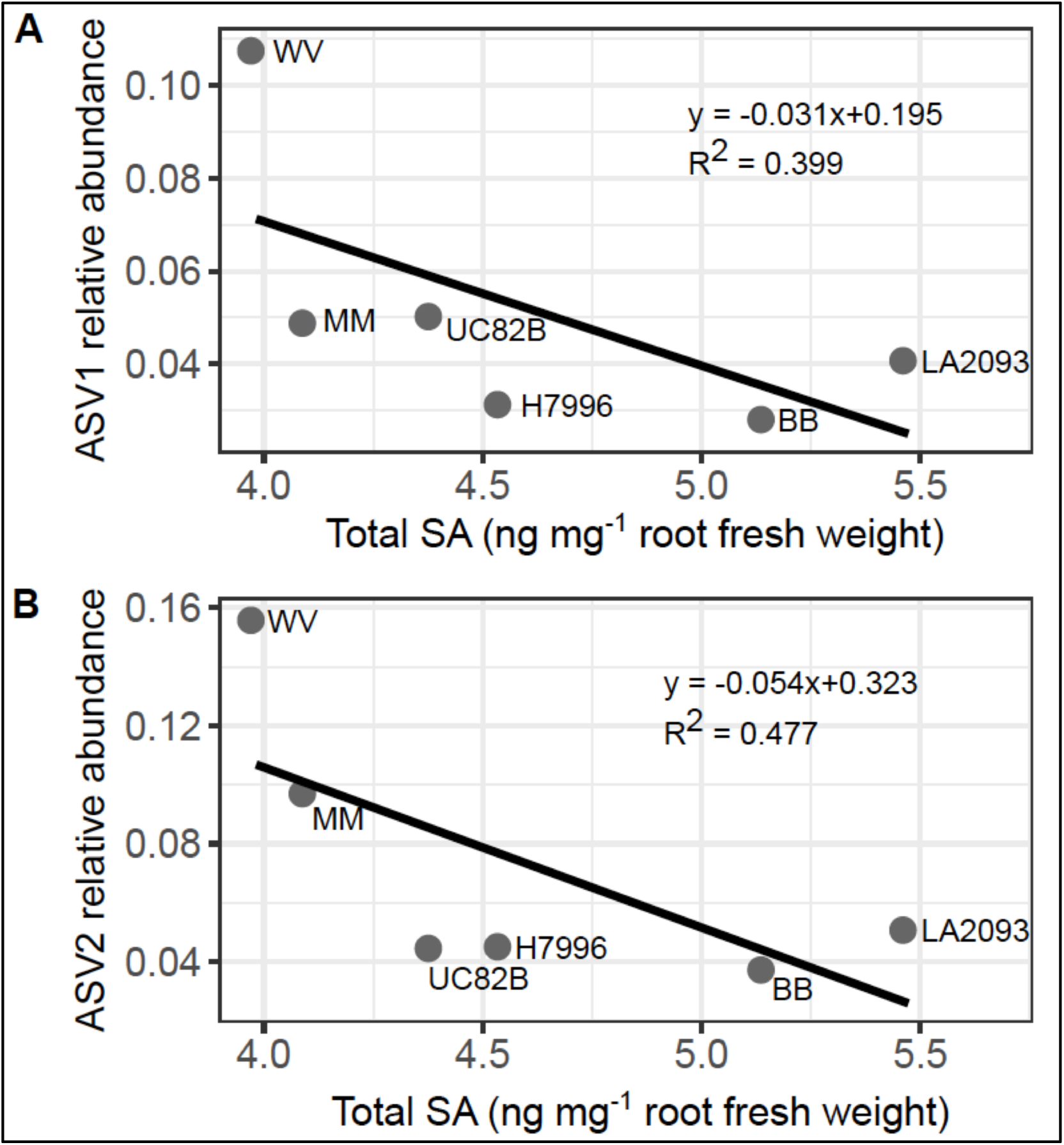
The relative abundance of ASV1/2 show a negative association with total SA content. Scatterplots with fitted trendlines comparing the average relative abundance of a) ASV1 (*Anaerobacillus*) and b) ASV2 (*Delftia*) versus total SA content in nanograms (ng) per milligram (mg) fresh weight (FW) of frozen, ground roots from six wild-type genotypes – UC82B, MM, H7996, WV, BB, LA2093 (n = 3 or 4 per genotype) grown in potting mix mixed with field soil. Points are labeled by genotype. Equations and R^2^ values for each trendline represented on each plot.

## DISCUSSION

Our results show that ET and SA hormone pathways contribute to the composition and diversity within the tomato root endosphere. In contrast to the root endosphere, the rhizosphere microbial community appears to be less sensitive to hormone deficiencies, as only the rhizosphere microbiome of the *ACD* mutant differed significantly from the wild-type background. The *NahG* and *ACD* mutants have decreased root endosphere α-diversity but increased abundance of two taxa (ASV1/2), a correlation that was also observed among 24 wild-type genotypes. In six wild-type genotypes, lower root SA levels were correlated with higher levels of ASV1/2. Based on these data, we propose that in tomato, the ET and SA pathways contribute to modulating the abundance of specific bacterial taxa in the root endosphere.

Whether the abundance of ASV1/2 directly impacts root endosphere α-diversity, and whether the ET and SA pathways directly impact diversity by altering growth of specific taxa (like ASV1/2) is not yet clear. Our data are consistent with the hypothesis that the plant immune system acts to maintain microbial homeostasis and controls the microbial load of commensal microbes [3].

### Species-specific effects of plant hormone pathways in the microbiome

We observed that the root endosphere and rhizosphere of the *ACD* transgenic had reduced α-diversity compared to its wild-type background. In another Solanaceous species, *Nicotiana attenuata,* mutants deficient in ET biosynthesis and signaling showed a small reduction in culturable bacterial diversity in the root endosphere compared to wild-type *N. attenuata* (Long *et al.* 2010) similar to our observation of reduced α-diversity in the tomato endosphere of *ACD*. We did not observe an effect of *Nr,* an ET perception and signaling mutant, on root microbiome assembly. It is important to note that *ACD* constitutively degrades ACC, the precursor to ET. Because the transgenic has decreased levels of both ACC and ET (Lanahan 1994), it is not clear whether the lowered α-diversity is due to a reduction in ACC, ET, or both. Alternatively, other effects of ACC degradation, including increased ammonium and alpha ketobutyrate, and potential impacts on the upstream pool of *S*-adenosyl-L-methionine (SAM), could lead to changes that result in lowered root endosphere α-diversity in *ACD.* In Arabidopsis, the Bray-Curtis index of a leaf epiphytic synthetic community increased in the ET signaling mutant *ein2* compared to wild-type plants (Bodenhausen, Bortfeld-Miller, Ackermann & Vorholt 2014). Thus, the effect of ET on the microbiome may differ between plant species or organ (root vs. leaf).

Similar to our observations for ET, we observed that the *NahG* mutant had decreased α-diversity. Because the *NahG* transgenic degrades SA to catechol, it is possible that increased catechol, and not decreased SA, led to the reduction in α-diversity. In Arabidopsis, increased SA levels may lead to a reduction in α-diversity in some compartments of the plant microbiome (Kniskern, Traw & Bergelson 2007; Lebeis *et al.* 2015). However, decreased SA (as in the SA biosynthesis mutant *sid2*) does not appear to have an effect on the α-diversity of the Arabidopsis root (Lebeis *et al.* 2015) or leaf microbiome (Kniskern *et al.* 2007). Thus, there may be species-specific differences in the role of the SA pathway in modulating the microbiome.

We did not find evidence for a role for the JA pathway in tomato root microbiome diversity. In Arabidopsis and wheat, JA functions in community composition and diversity (Carvalhais *et al.* 2013, 2015; Liu *et al.* 2017). Like the ET and SA pathways, the effects of the JA pathway on the root microbiome may be species-specific.

There appeared to be little impact of tomato hormone deficiencies on rhizosphere diversity, as only one mutant (*ACD*) was significantly different than its wild-type background. This may have been due to the young age of our plants, as plant age plays a role in determining the effects on the root microbiome (Wagner *et al.* 2016).

### Natural variation in the α-diversity of wild-type root endosphere microbiomes negatively correlates with colonization of specific ASVs

The root endosphere of *NahG/ACD* plants were enriched for two taxa (ASVs 1 and 2) that were classified into genus *Anaerobacillus* (Phylum: Firmicutes) and *Delftia* (Phylum: Proteobacteria), respectively. *Delftia* is a highly diverse root-colonizer, and some isolates have been characterized for their biocontrol and plant growth promotion (PGP) properties (Morel, Cagide, Minteguiaga, Dardanelli & Castro-Sowinski 2015; Agafonova *et al.* 2017).

*Anaerobacillus* is not a well-characterized genus (Zavarzina, Tourova, Kolganova, Boulygina & Zhilina 2009), but this ASV is also identical to some isolates from the *Bacillus* genus, which is well known for its biocontrol and PGP properties (Santoyo, Orozco-Mosqueda & Govindappa 2012; Santoyo, Moreno-Hagelsieb, del Carmen Orozco-Mosqueda & Glick 2016; Takahashi *et al.* 2014; Cao *et al.* 2018).

The correlation between enrichment of these taxa and lower α-diversity in the root endospheres of *NahG* and *ACD* did not appear to be due solely to the effects of *NahG* and *ACD* degradation products, because we observed a similar correlation among a panel of wild-type tomatoes. Further, SA levels of wild-type tomato roots were moderately negatively correlated with relative abundance of ASV1/2. Whether ASV1/2 ‘outcompete’ other taxa in low SA and low ET environments will be tested in future work.

### Tomato genotype impacts Shannon diversity, richness and composition in the tomato root endosphere but the rhizosphere is less impacted by genotype

In our experiments, diversity of the root endosphere varied among all wild-type tomato genotypes (20 RILs, H7996, WV, LA2093, BB, MM, UC82B, Pearson, Castlemart; Figure 2, Figure 5c and 5d). In contrast, the rhizosphere microbiome appeared more stable to variations in genotype (Figure 2). Genotypes included both fresh market and processing cultivated tomatoes (*S. lycopersicum*) and *S. pimpinellifolium,* the closest wild relative to *S. lycopersicum*. WV, an *S. pimpinellifolium* that is highly susceptible to the soil borne bacterial pathogen *Ralstonia solanacearum,* had lower α-diversity in the root endosphere compared to H7996, an *S. lycopersicum* genotype resistant to *R. solanacearum* (Table S18). However, α-diversity appeared to be unrelated to resistance because the endosphere α-diversity of cultivar BB, which is also susceptible to *R. solanacearum,* did not significantly differ from resistant lines H7996 and LA2093. Our results suggest that tomatoes are more selective about which taxa colonize the root endosphere, and even genetic differences within species may alter root endosphere colonization. Further work will be required to discover the ecological costs or benefits that may be associated with differences in root community α-diversity.

In conclusion, our results reveal roles for the ET and SA pathways in α-diversity of the tomato root endophytic microbiome. Further, our data suggests a genetic basis for colonization of specific taxa. These findings are important both for our basic understanding of how plants regulate microbial colonization by non-pathogenic microbes, and suggest that breeding tomatoes for improved associations with root microbe communities is possible.

## ACKNOWLEDGEMENTS

We thank the Purdue Genomics Core for helpful discussions, Gregg Howe for *def1* and CMII seeds, Harry Klee for *Nr* and Pearson seeds, and members of the Iyer-Pascuzzi lab for helpful comments. This work was funded by Purdue University start-up funds, Purdue Center for Plant Biology seed grant funds, Hatch funds (project numbers IND011293 to AIP and 177845 to JRW), and a grant from the Foundation for Food and Agriculture Research (FFAR) to AIP. This material is based upon work supported by the National Science Foundation Graduate Research Fellowship Program under Grant No. DGE-1333468 to EF. Any opinions, findings, and conclusions or recommendations expressed in this material are those of the authors and do not necessarily reflect the views of the National Science Foundation.

## AUTHOR CONTRIBUTION

EF, JW, and AIP designed experiments; EF, MJ, performed experiments and data analysis; EF, MJ, JW and AIP wrote the paper.

Sequences and metadata have been deposited into the NCBI BioProject database with BioProjectID PRJNA507021 (http://www.ncbi.nlm.nih.gov/bioproject/507021).

Code used for microbiome analysis is available at the Purdue University GitHub (Iyer-Pascuzzi group): https://github.rcac.purdue.edu/AnjaliIyerpascuzziGroup/Tomato-Root-16S-Sequencing.

## Supporting Information

**Figure S1** Overall patterns of β-diversity across all samples (hormone mutants and their wild-type backgrounds, RILs and parents) cluster by compartment.

**Figure S2** *ACD* and *NahG* differ from their respective wild-types in β-diversity.

**Figure S3** Phyla and relative abundance of endosphere-enriched ASVs (top) and endosphere-depleted ASVs (bottom) in mutants and their respective wild-type backgrounds.

**Figure S4** Taxonomy of shared and unique ASVs between *NahG* and *ACD*.

**Table S1** Soil characteristics

**Table S2** Mutant lines and backgrounds

**Table S3** *S. lycopersicum, S. pimpinellifolium* lines and RILs

**Table S4** Sequencing summary

**Table S5** Raw count table of all 901 ASVs across all samples

**Table S6** Sequencing sample metadata

**Table S7** Taxonomy table

**Table S8** Summary of relative abundance of taxa averaged at family level in hormone mutants and their respective wild-type backgrounds

**Table S9** Summary of differential abundance from rhizosphere to endosphere by genotype

**Table S10** Full differential abundance results for *ACD*

**Table S11** Full differential abundance results for UC82B

**Table S12** Full differential abundance results for *NahG*

**Table S13** Full differential abundance results for MM

**Table S14** Full differential abundance results for *Nr*

**Table S15** Full differential abundance results for Pearson

**Table S16** Full differential abundance results for *def1*

**Table S17** Full differential abundance results for CMII

**Table S18** Average α-diversity in endosphere and rhizospheres of wild-type tomato genotypes

