## Supplementary material for "Defense hormones modulate root microbiome diversity and composition in tomato": Figure S1

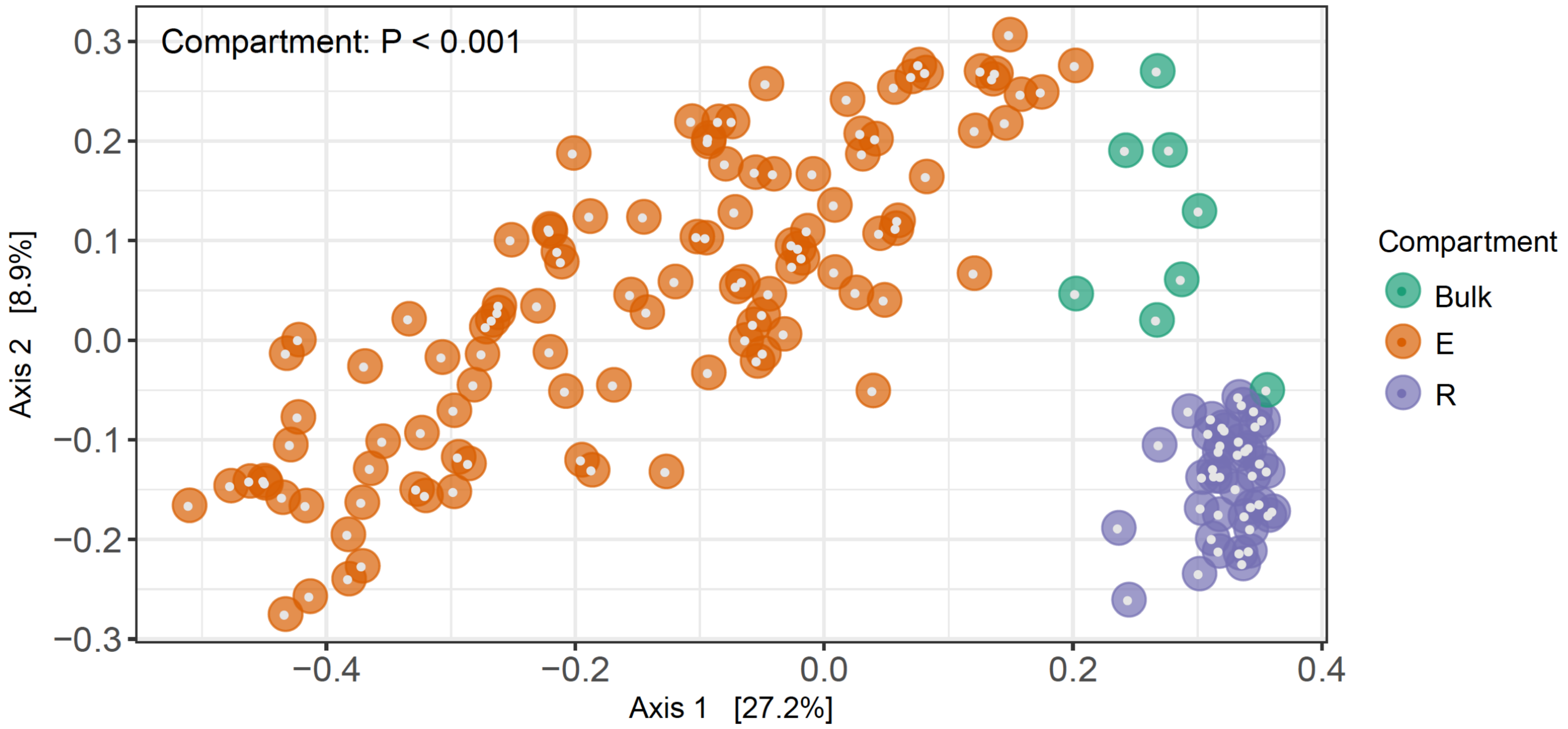


**Figure S1** Overall patterns of beta diversity across all samples (hormone mutants and their wild-type backgrounds, RILs and parents) cluster by compartment.
