## Supplementary figures and images for "Defense hormones modulate root microbiome diversity and composition in tomato"

### Figure S2

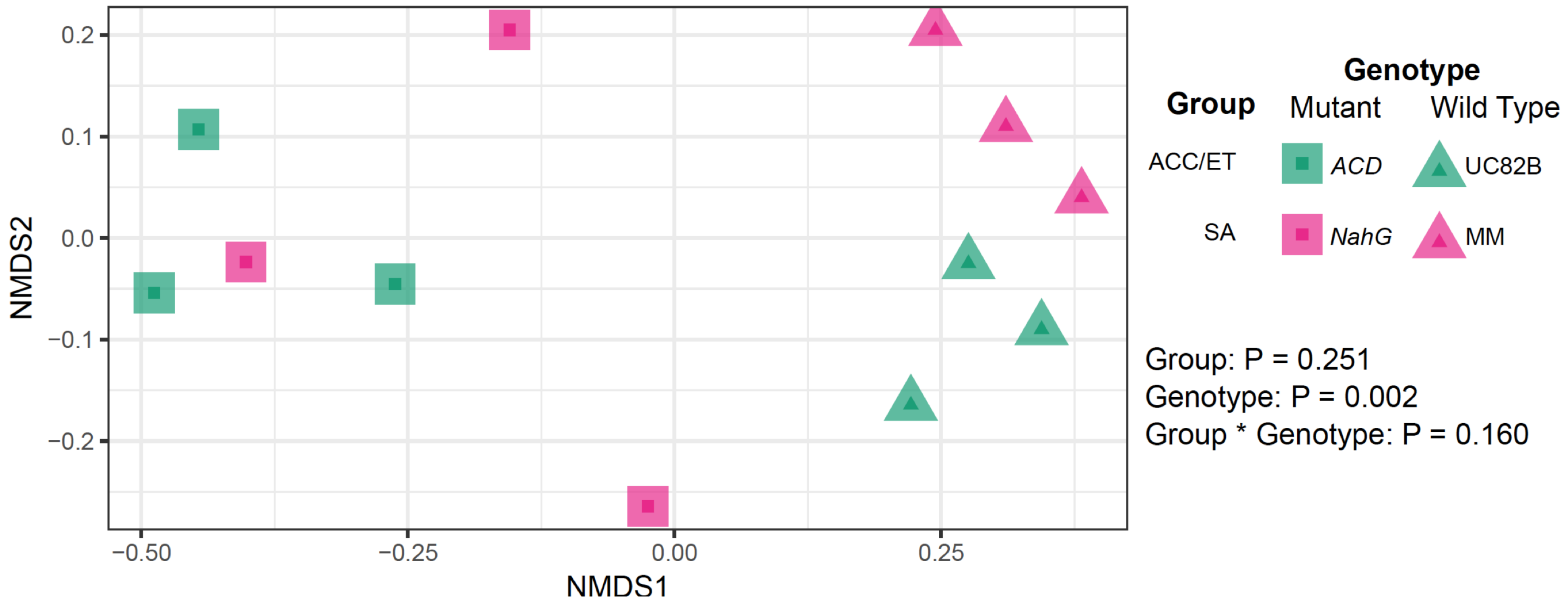


**Figure S2** *ACD* and *NahG* differ from their respective wild-types in beta diversity

### Figure S4

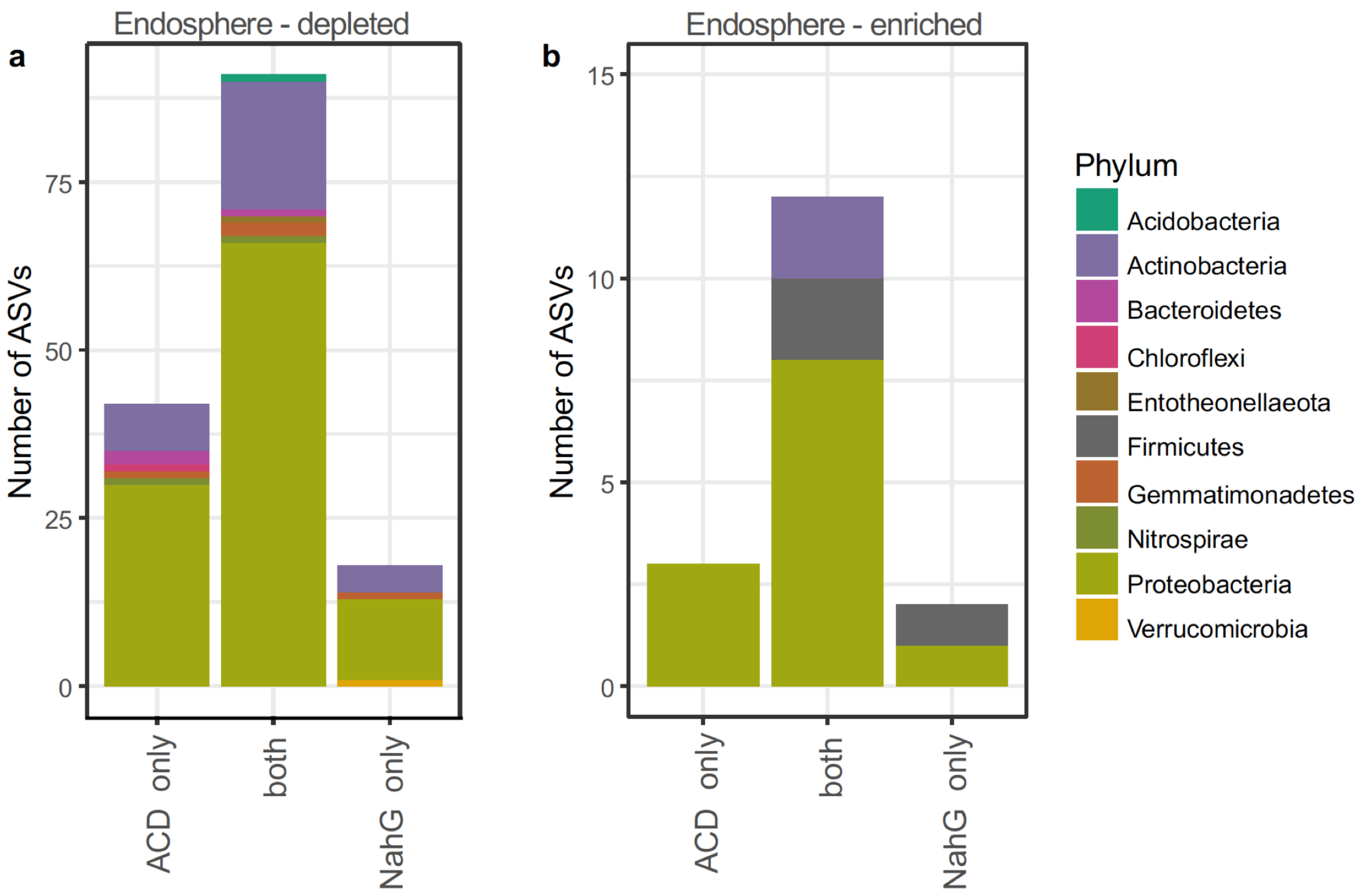


**Figure S4** Taxonomy of shared and unique ASVs between *NahG* and *ACD*.
