## Supplementary material for "Defense hormones modulate root microbiome diversity and composition in tomato": Figure S3

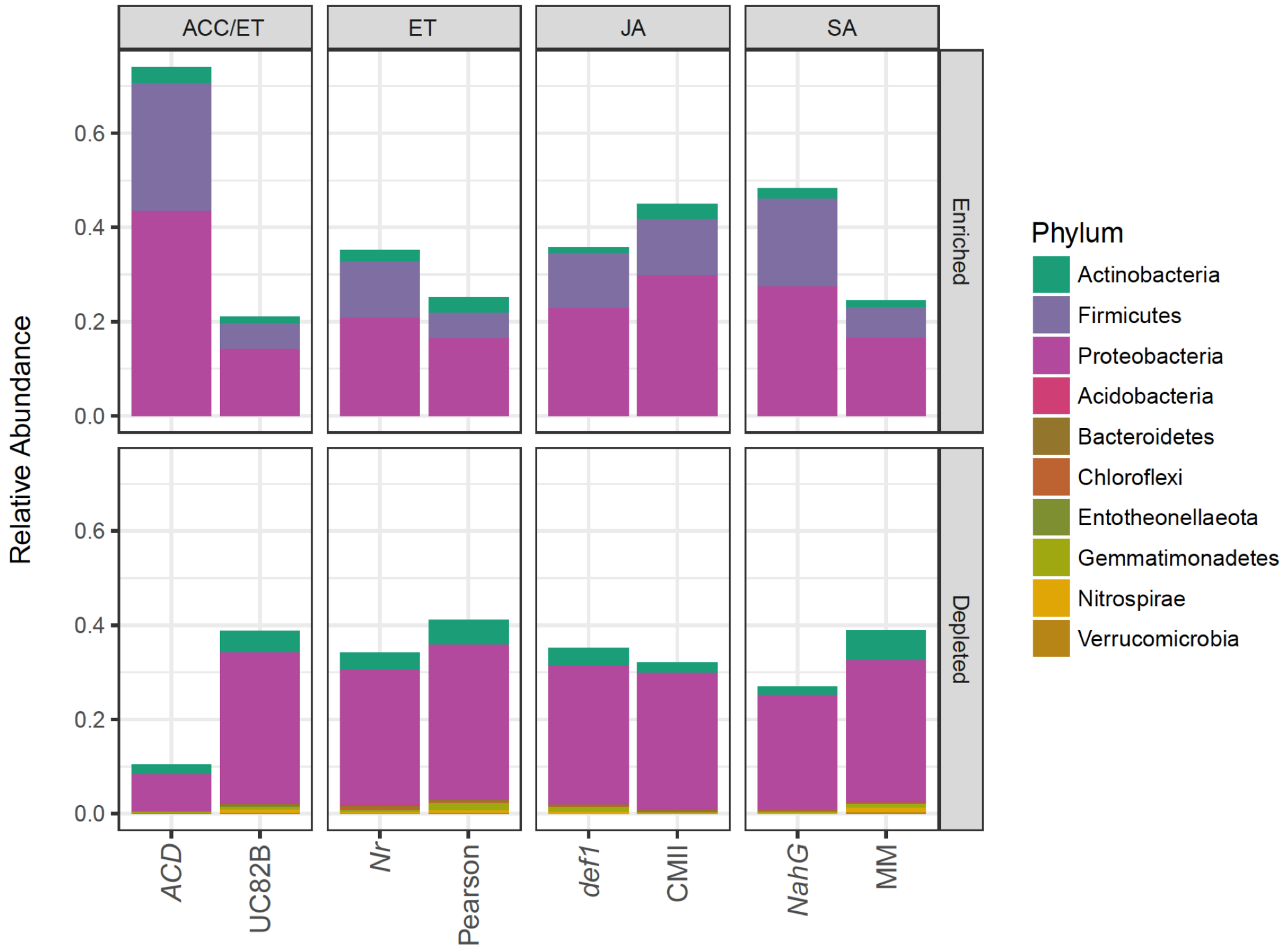


**Figure S3** Phyla and relative abundance of endosphere-enriched ASVs (top) and endosphere-depleted ASVs (bottom) in mutants and their respective wild-type backgrounds.
